# The Type 9 Secretion System enables sharing of fungal mannan by human gut *Bacteroides*

**DOI:** 10.1101/2022.07.15.500217

**Authors:** Ekaterina Buzun, Tiaan Heunis, Curtis Cottam, Carl Morland, Matthias Trost, Elisabeth C Lowe

**Affiliations:** Biosciences Institute, Faculty of Medical Sciences, Newcastle University, Newcastle upon Tyne, NE2 4HH, UK; Department of Pathology, University of California San Diego, La Jolla, CA, United States

## Abstract

Degradation of complex carbohydrates in the gut is a key trait of *Bacteroides* species. Some glycans are metabolised ‘selfishly’ releasing few or no oligosaccharide breakdown products from complex polysaccharides, whereas others release oligosaccharides and cross feed other microbes. The outer cell wall of many fungi commonly found in the gut consists of highly α-mannosylated proteins which have been shown to be metabolised in a ‘selfish’ manner by *Bacteroides thetaiotaomicron*. We show that the species *Bacteroides salyersiae* releases branched manno-oligosaccharides during growth on mannan and that these act as a nutrient source for *Bacteroides* spp. that are unable to degrade polymeric mannan. Molecular characterisation of the locus responsible for mannan degradation reveals that it contains multiple glycoside hydrolases and glycan binding proteins targeted to the Type 9 Secretion System, a Bacteroidetes specific secretion system that allows the secretion of large folded proteins across the outer membrane. More commonly found in oral and environmental Bacteroidetes, here the T9SS enables *B. salyersiae* to locate large, multimodular enzymes and glycan binding proteins outside the cell to target a complex, branched polysaccharide. This points to a previously unknown role of the T9SS in glycan metabolism in gut *Bacteroides*.

## Introduction

Gram-negative bacteria possess several secretion systems (I-X) enabling them to locate proteins to the cell surface, the extracellular environment, or directly into other cells (1, 2). The Type 9 Secretion System (T9SS) is restricted to species within the Bacteroidetes phylum. Best characterised in the oral microbe, *Porphyromonas gingivalis* (3, 4), and the environmental bacterium, *Flavobacterium johnsoniae* (5, 6), the T9SS is a multiprotein complex which recognises a C-terminal signal domain (CTD) of periplasmic proteins, directing them to the surface, where a sortase conjugates them to a membrane molecule such as LPS in the case of *P. gingivalis* or secretes them into the environment in a soluble form (7–13). The proteins secreted via T9SS include proteases, which serve as virulence factors in *P. gingivalis* (3, 14), glycoside hydrolases (GH) (15), and proteins mediating gliding motility in *F. johnsoniae* (16–18).

Though the T9SS is common among Bacteroidetes, it was thought to be largely absent from the numerous anaerobic *Bacteroides* species, which are major colonisers of the human gut (6). The *Bacteroides* are of interest due to their role in degradation of the complex carbohydrates they encounter in the gut (19), often possessing hundreds of carbohydrate active enzymes (20, 21) specific for glycans from dietary, host and microbial sources (22–29). These enzymes are typically found within co-regulated loci called PULs (polysaccharide utilisation loci), alongside binding proteins, glycan transporters and sensor regulators (30, 31). A single glycan can be deconstructed by one or more PULs, depending on the complexity of the structure of the target polysaccharide (32, 33). Typically, PUL-encoded enzymes and glycan-binding proteins are directed to the surface as lipoproteins where they are attached to membrane lipids through a conserved cysteine residue (34–36). Type 9 secreted proteins have not been described to be encoded in PULs to date.

*Bacteroides* species utilise complex carbohydrates in a preferential order and often deploy either a ‘selfish’ or ‘sharing’ strategy for their degradation (37–40). It was previously shown that a common gut bacterium, *Bacteroides thetaiotaomicron* VPI-5482, degrades a major component of the fungal cell wall, α-mannan, via a ‘selfish’ mechanism (41). Fungal α-mannan shields cell wall proteins and underlying immunogenic glucans (**Fig. 1a**). O-linked mannan is short and linear, whereas N-linked is composed of an α-1,6-linked backbone which is heavily decorated with sidechains, extended by α-1,2/ α-1,3-mannosidic bonds (**Fig. 1b**) (42). Some of the side sidechains can be further decorated with additional short α-1,3 linked mannophosphate branches, which are attached to the structure via phosphodiester bonds (**Fig. 1b**) (42). *B. thetaiotaomicron* dedicates three α-mannan specific PULs encoding a range of enzymes and transport proteins to deconstruct this highly complex polysaccharide (41). Elements of these PULs were also found in *B. xylanisolvens* NLAE-zl-p352 and *B. ovatus* ATCC 8483, which both displayed poor growth on yeast mannan (41). Here we show that another common gut bacterium, *Bacteroides salyersiae* DSM 18765/WAL10018, deploys a ‘sharing’ mechanism for yeast mannan utilisation. The PUL orchestrating α-mannan breakdown in *B. salyersiae* does not share structural synteny with α-mannan PULs in *B. thetaiotaomicron* and contains enzymes and binding proteins which are directed for secretion via the T9SS. These proteins generate a pool of extracellular branched mannooligosaccharides, which are then utilised as ‘public goods’ by non-mannan degrading gut *Bacteroides*.

**Figure 1.**
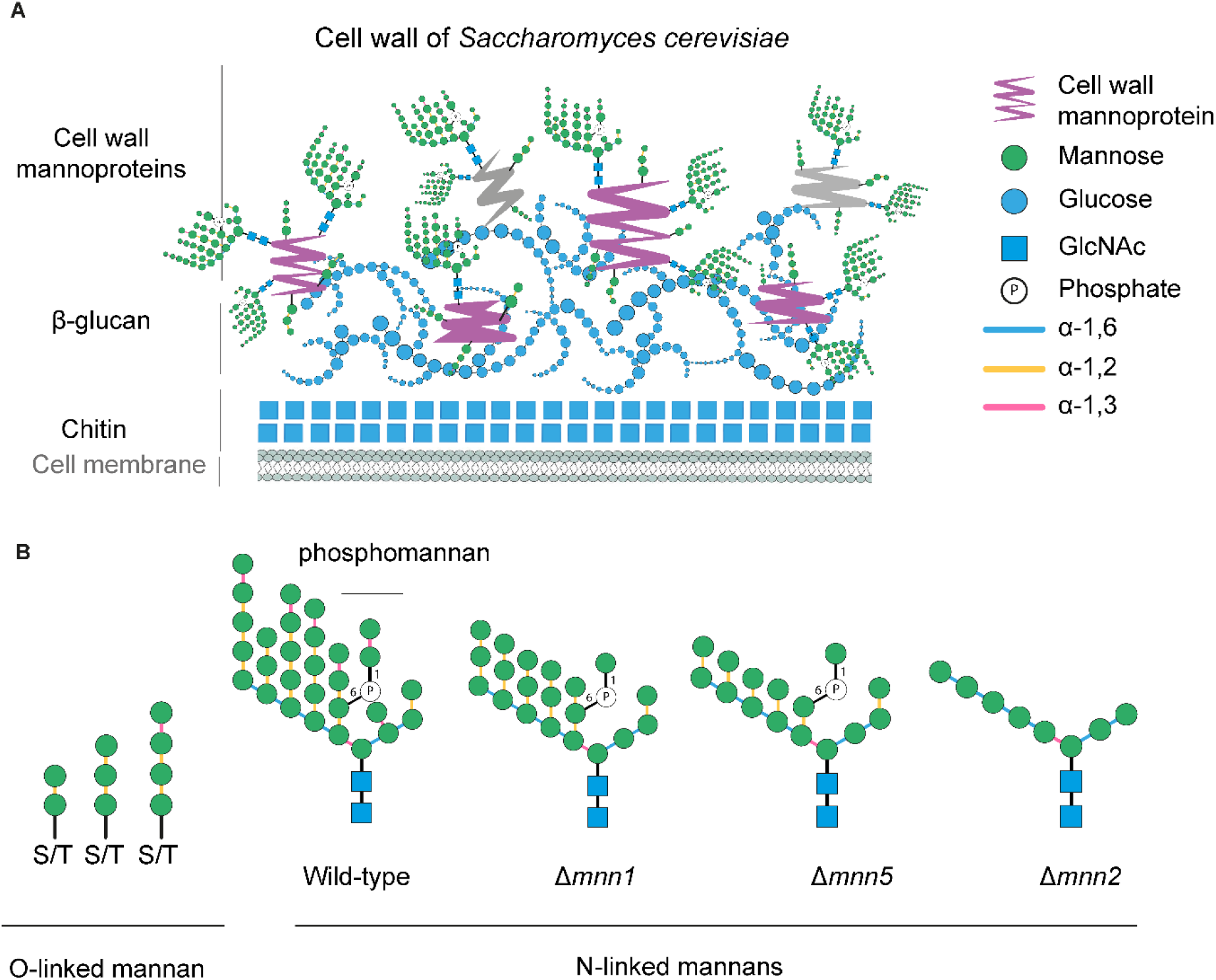
Organisation of the cell wall from *Saccharomyces cerevisiae*. a) layered structure of the cell wall, inner chitin and β-glucan layers are shown in blue, mannan is shown in green. b) types of mannosylation present in yeast cell wall. O-linked glycans form short linear α-1,2/α-1,3 linked glycans. N-mannan from wild type *S. cerevisiae* possesses a branched structure, where α-1,6-linked backbone is highly decorated with α-1,2/α-1,3-linked sidechains. Phosphomannan is α-1,3 linked. S. cerevisiae Δmnn1 strain lacks α-1,3-decorations due to the absence of α-1,3-mannosyltransferase (MNN1); mnn5 is deficient in α-1,2-mannosyltransferase resulting in the α-1,6-backbone with a single α-1,2-linked mannose attached; mannan from Δmnn2 strain is composed of a linear α-1,6-linked backbone due to the lack of enzymes necessary for side chain biogenesis.

## Results

### *B. salyersiae* mediates yeast mannan breakdown at the cell surface

We first assessed the ability of 19 different *Bacteroides* species to grow on mannan from *Saccharomyces cerevisiae*. In line with previous data (41), both *B. thetaiotaomicron* VPI-5482 and *B. salyersiae* WAL 10018 displayed robust growth on yeast mannan (**Fig. 2a**). *B. salyersiae* has a 20-hour lag phase but reached a higher maximum OD_600_ than *B. thetaiotaomicron* (**Fig. 2a**). Analysis of the cell free supernatant post bacterial growth revealed that in contrast to *B. thetaiotaomicron, B. salyersiae* accumulated oligosaccharides in the media (**Fig. 2b**). We analysed the constituent glycosidic bonds of these oligosaccharides, using previously characterised GH92 α-mannan specific enzymes from *B. thetaiotaomicron*, which can remove α-1,3 (BT3862 and BT3858) and α-1,2 (BT4092) mannosyl decorations from mannan side chains (43). We also used a GH125 α-1,6-mannosidase, BT2632, capable of deconstructing the α-1,6-linked mannan backbone (41). This exo-mannosidase degradation assay revealed that the extracellular oligosaccharides were branched and retained α-1,3 and α-1,2 side chain decorations (**Fig. S1**). To confirm that the oligosaccharides were generated at the cell surface and were not a result of cell lysis, we assessed activity of the whole cells, using Proteinase K to remove surface enzymes. This showed that *B. salyersiae* gradually releases oligosaccharides and mannose from yeast mannan using enzymes located on its cell surface (**Fig. 2c**).

**Figure 2.**
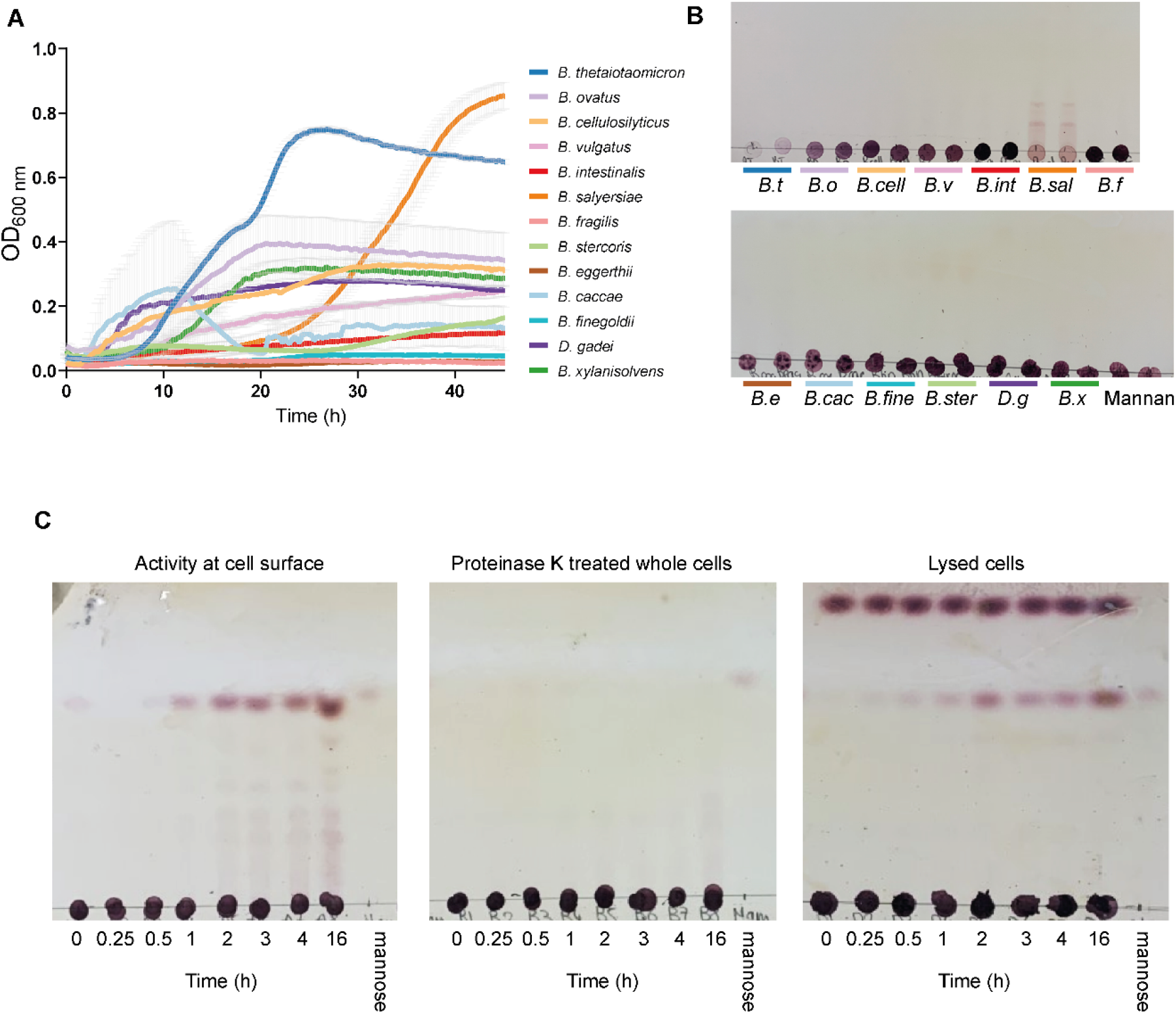
*Bacteroides salyersiae* releases manno-oligosaccharides during growth on fungal α-mannan. a) Growth of selected Bacteroidetes strains on defined media containing *S. cerevisiae* mannan. b) TLC of growth media from stationary phase of the cultures in panel a. c) Enzyme activity assessed by TLC from whole *B. salyersiae* cells grown on yeast mannan (left), whole cells after treatment with proteinase K (middle) and lysed cells (right).

### Identification of *B. salyersiae* mannan degrading apparatus

We carried out whole cell proteomics on mid-log phase glucose or α-mannan grown *B. salyersiae* and quantified 2188 proteins (**Fig. 3a, Table S1**), 58 of which were significantly upregulated (p<0.05,>2-fold change) during growth on α-mannan. These were grouped into 3 distinct loci (**Fig. 3a and c**). For brevity, locus tags in *B. salyersiae* are named HMPREF1532_XXXXX which will be replaced with BsXXXXX. Genome analysis showed that *B. salyersiae* possesses homologs of the T9SS structural proteins (**Fig. 3b**), which shared low similarity with the structural proteins from *P. gingivalis* or *F. johnsoniae* (**Table S2**). From these, all of the essential components of the system, including the main translocon SprA (44) (Bs03761) and the central sorting protease PorU (9) (Bs04019), were not upregulated but present in the proteome profile of *B. salyersiae* during growth on both mannan and glucose (**Fig. 3b**). The two most highly upregulated loci are depicted in **Fig. 3c**. One locus lacked a pair of SusC/D proteins and is therefore described as a CAZyme cluster (**Fig. 3c**). The largest locus is identified as ‘predicted PUL52’ in the *B. salyersiae* genome in the database PUL-DB (45), which encloses genes Bs04071 to Bs04080. These proteomics data suggest the PUL extends as far as Bs04092, including a further six GH enzymes and two sensor-regulators. GH families in this large PUL include predicted α-mannanases (GH76) and α-mannosidases (GH38, GH92, GH125) as well as GH2, GH3 (broad specificity β-glycosidases) and GH97 (α-galactosidase or glucosidase). As previously characterised in *P. gingivalis* and *F. johnsoniae*, Type A substrates of the T9SS possess conserved C-terminal amino acid motifs residing in the last 80 amino acids, identified in the TIGRfam database as Por_Secre_tail, TIGR04183 (46, 47). Alignment of the terminal tails of the proteins in this PUL revealed that at least 10 proteins contain T9SS associated motifs, including five GH enzymes (**Fig. 3c and Fig. S2**). This includes a GH38, Bs04085, and a GH76, Bs04077. The GH76 is much larger than the surface GH76 endo-mannanases characterised from *B. thetaiotaomicron*, and contains a carbohydrate binding module; CBM6, a ricin-lectin domain and several other domains in addition to the catalytic domain (**Fig. 3d**). Further analysis of *Bacteroides* genomes for the Por_Secre_tail domain showed that *B. salyersiae* likely directs at least 99 proteins for secretion by the T9SS, including multiple glycoside hydrolases (**Table S3**), indicating the T9SS plays an important role in the glycan degrading capacity of *B. salyersiae*. In addition to *B. salyersiae*, we found likely T9SS substrates in strains of *B. nordii*, *B. cellulosilyticus*, and *B. intestinalis*.

**Figure 3.**
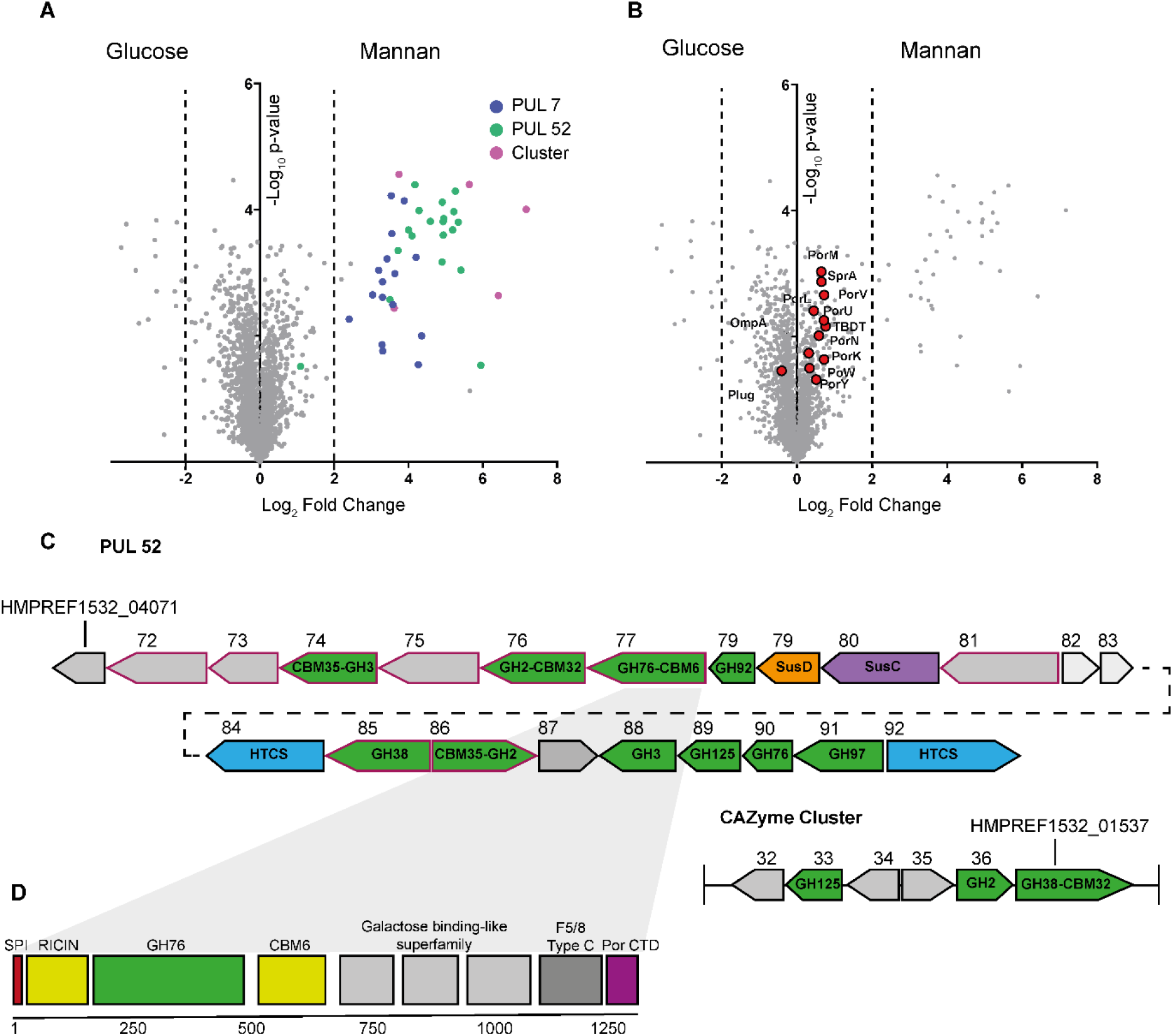
Comparative proteomics of mannan-grown *B. salyersiae*. Whole cell proteomic analysis of glucose and α-mannan grown cells (triplicate growths) harvested at midexponential phase. a) Mannan vs glucose fold-change. The upregulated loci include PUL7 (blue), PUL52 (green) and CAZyme cluster (pink). b) Mannose vs glucose fold change, highlighting components of the T9SS which were detected in both glucose and mannan grown cells. c) Architecture of PUL52 and CAZyme Cluster. CAZymes are coloured green, SusC/D pair in purple and orange, and HTCS (hybrid two-component system sensor regulators) in blue. ORFs of unknown function are coloured grey. Proteins predicted to encode T9SS signal CTDs are outlined in purple. d) Domain architecture of Bs04077^GH76-CBM6^. Catalytic domain is green, potential lectin or carbohydrate binding domains in yellow, T9SS C-terminal signal sequence in purple, and other domains in grey. The periplasmic SPI signal sequence is red.

### Yeast mannan degradation in *B. salyersiae*

The α-1,2/α-1,3-linked sidechains sterically restrict the α-1,6 linked backbone from the enzymatic degradation by endo-acting mannanases from the GH76 family (**Fig. 1b**). In *B. thetaiotaomicron*, mannan breakdown is initiated by three endo-acting enzymes, GH99 (BT3862) and two GH76 (BT3792 and BT2623) located at the cell surface, whereas essential exo-acting enzymes from the GH92, GH38, and GH125 families are primarily in the periplasm (41). In *B. thetaiotaomicron*, a periplasmic GH38, BT3774, acts as a central broad-specificity α-mannosidase capable of removing α-1,3; α-1,2, and mannose-1-phosphate decorations, exposing the backbone to GH76 mannanases and GH125 mannosidases (41).

*B. salyersiae* upregulates two GH38s: Bs04085 and Bs01537 (**Fig. 3c**). While Bs01537 is 70% identical to BT3774, Bs04085 shares low similarity with either BT3774, Bs01537 or other characterised GH38s (**Fig. S3A**). Both Bs04085 and Bs01537 retain the N-terminal SP1 signal peptide, indicating that both enzymes are periplasmic. Our alignment showed that the C-terminus of Bs04085 but not Bs01537 contains T9SS associated amino acid motifs (**Fig. S2**), suggesting that this enzyme could be further translocated to the cell surface or extracellular milieu via the T9SS. We characterised Bs04085 and Bs01537 with a variety of biochemical assays and showed that both GH38s act as broad-specificity α-mannosidases and are capable of cleaving mannose-1-phosphate linkages (**Fig. S3**). Unlike Bs01537, Bs04085 was able to efficiently hydrolyse highly branched α-mannan, which retains both α-1,3 and α-1,2 sidechain decorations (**Fig. 4a**). In contrast, Bs01537 displays a preference for small oligosaccharides and less complex mannan polymers isolated from *S. cerevisiae* glycosyl-transferase mutants (42), which only contain α-1,2-mannosyl decorations (**Fig. 4a, Fig. S3**). The GH92 Bs04078 is active against α-1,2 and α-1,6 mannobiose.

**Figure 4.**
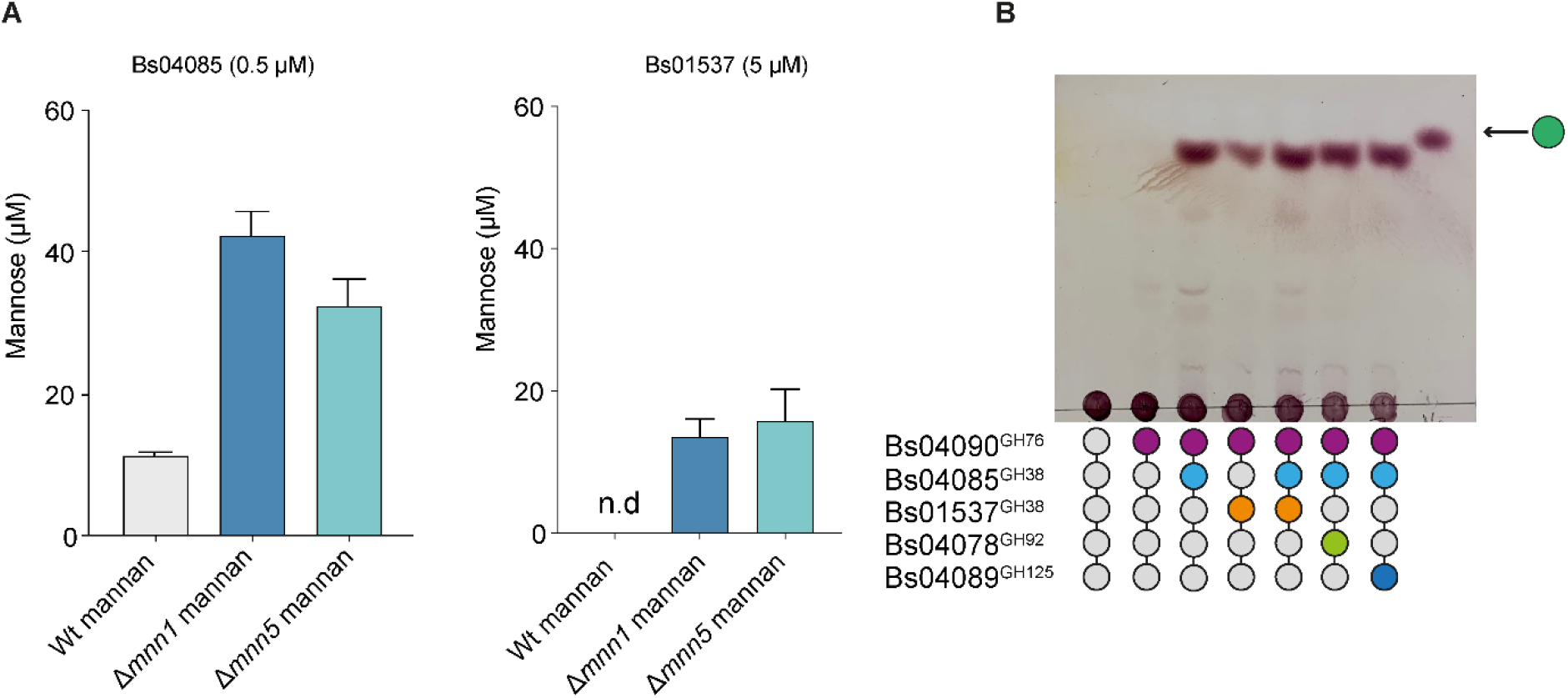
Activity of recombinant enzymes from *B. salyersiae* against yeast α-mannan. a) Total amount of mannose released from wild type, Δmnn1, Δmnn5 *S. cerevisiae* strains by Bs04085^GH38^ and BS01537^GH38^. n.d - none detected. b) Activity of recombinant enzymes from *B. salyersiae* PUL52 against yeast mannan.

In addition to GH38s, two GH76s (Bs04077 and Bs04090) were upregulated in *B. salyersiae* during growth on α-mannan. As described above, Bs04077 is a likely substrate of the T9SS. Bs04090 possesses a lipoprotein signal sequence which suggests it could also be located on the outer membrane. Despite extensive attempts with multiple cloning vectors, *E. coli* strains and gene synthesis services, cloning of full length Bs04077 or its catalytic domain into an expression vector failed. All attempts resulted in large deletions within the cloned gene, or insertion of *E. coli* genomic DNA fragments into the plasmid, suggesting toxicity of the gene or protein product in *E. coli*. Bs04090 displayed poor activity against wild type α-mannan (**Fig. 4b**) and its mannanase activity was enhanced by GH38 Bs04085 which likely exposes the a-mannan backbone, allowing release of some oligosaccharides (**Fig. 4b**). Consistent with its lack of activity against WT mannan, addition of GH38 Bs01537 did not enable Bs04090 to access the mannan backbone (**Fig. 4b**). Addition of other recombinant mannosidases from the PUL: GH92 Bs04079 and GH125 Bs04089 resolved the products of the GH76/GH38 enzymatic cocktail (**Fig. 4b**), however the products of theses reactions were different to the oligosaccharides generated by the whole cells (**Fig. 2c**). This suggests that mannan degradation at the cell surface involves an additional endo-acting enzyme, likely Bs04077. Bs04085’s T9SS domain predicts it is located on the cell surface of *B. salyersiae*, where it acts as a broad-spectrum exo-mannosidase, generating extracellular mannose observed in the whole cell assays (**Fig. 2c**). Bs01537, similar to its homologue in *B. thetaiotaomicron*, orchestrates depolymerisation of shorter and less complex manno-oligosaccharides in the periplasm.

### Type 9 secretion of yeast α-mannan binding proteins

The T9SS mediates extracellular secretion of a chitinase, ChiA, in *F. johnsoniae* (15). Although gut *Bacteroides* are not known to secrete soluble glycoside hydrolases into the extracellular milieu, we performed a secretion assay to assess this. Cell-free culture supernatant was collected at multiple points during growth, concentrated 5-fold and examined with SDS-PAGE for the presence of proteins. This showed that proteins indeed accumulated in the culture supernatant (**Fig. 5a**), though no enzymatic activity was detected against any mannan substrates. In-gel MALDI-TOF analysis of the two most abundant proteins identified them as a 138 kDa protein of unknown function (Bs04081) and 76 kDa SusD-like protein (Bs04079), both of which are found in the *B. salyersiae* mannan PUL52 (**Fig. 5a**). SusD-like proteins usually associate with SusC-like proteins to mediate import of oligosaccharides (48, 49), therefore its presence in the extracellular space in a soluble form was unlikely. Species such as *B. thetaiotaomicron* and *B. fragilis* release SusD-like proteins, other putative binding proteins, and glycoside hydrolases in outer membrane vesicles (OMVs) (50–52). Transmission Electron Microscopy (TEM) of the pellet obtained from the ultracentrifuged supernatant showed that *B. salyersiae* produces OMVs of varying sizes during growth on yeast mannan and glucose (**Fig. 5b**). To further investigate whether these proteins were released with the OMVs or secreted in a soluble form, we concentrated ultracentrifuged supernatant and analysed it with SDS-PAGE alongside OMVs. This showed that both proteins were detected in the OMV fraction, not the supernatant, indicating that they associate with the OMVs rather than as soluble secreted proteins (**Fig. S4**)

**Figure 5.**
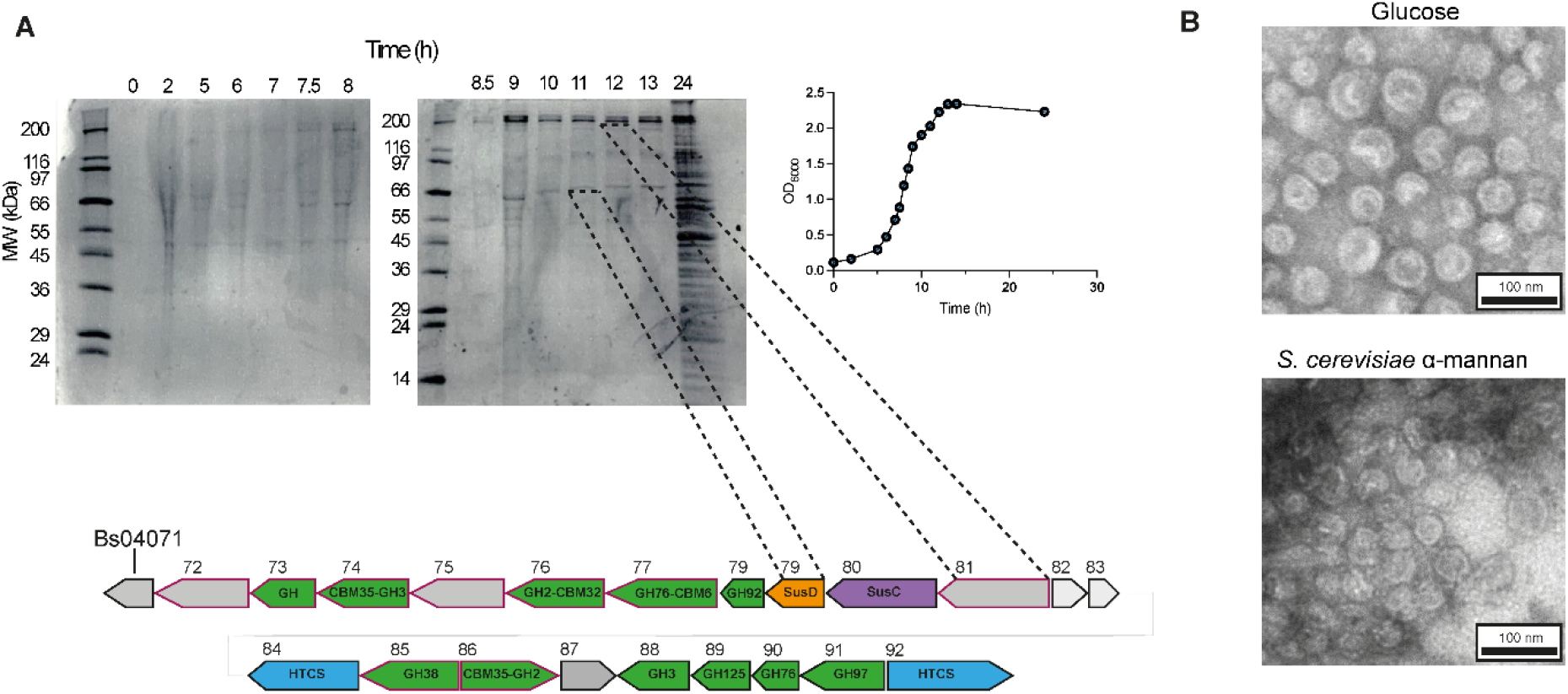
a) SDS-PAGE of cell-free supernatant collected during *B. salyersiae* growth on yeast mannan. Proteins from the two clearest bands were excised and analysed with MALDI-TOF. The largest band was identified as Bs04081, a 136 kDa protein from *B. salyersiae* PUL52, the second protein was identified as Bs04079, 80 kDa SusD-like protein from the same PUL. b) *B. salyersiae* produces outer membrane vesicles (OMVs) during growth on glucose and yeast mannan. OMVs were pelleted by ultracentrifugation and viewed with TEM using negative staining. Pictures were acquired at 20,000 x magnification at 100 kV accelerating voltage, 100 nm scale bar shown.

The domain organisation of Bs04081 includes a Big-2 domain (IPR003343, Pfam 02368) which is annotated as a potential adhesin, indicating that it may act as a binding protein (**Fig. 6a**). We also looked for the presence of such domains in other proteins within the main mannan PUL and identified another large protein (240 kDa) of unknown function, Bs04072, predicted to contain the Big-2 domain as well as multiple other domains (**Fig. 6b**). Neither protein showed any enzymatic activity against mannan substrates, but binding assays using affinity gels and Isothermal Titration Calorimetry demonstrated that Bs04081 indeed binds yeast mannan, displaying a preference for the α-1,2 decorated mannan variants over the undecorated α-1,6-linked backbone (**Fig. 6c and Fig. S4**). In contrast, Bs04072 exhibited weaker binding to branched yeast mannan. (**Fig. 6d,e**). Neither of the proteins bound to mannan from *Candida albicans*, where α-mannan is capped with additional β-1,2-mannosyl decorations (53) (data not shown), indicating that both proteins only recognise α-linked mannose units. Our alignment showed that Bs04072 and Bs04081 possess T9SS signals in their C-termini (**Fig S2**), suggesting a role for the T9SS in secretion of large multi-modular polysaccharide binding proteins.

**Figure 6.**
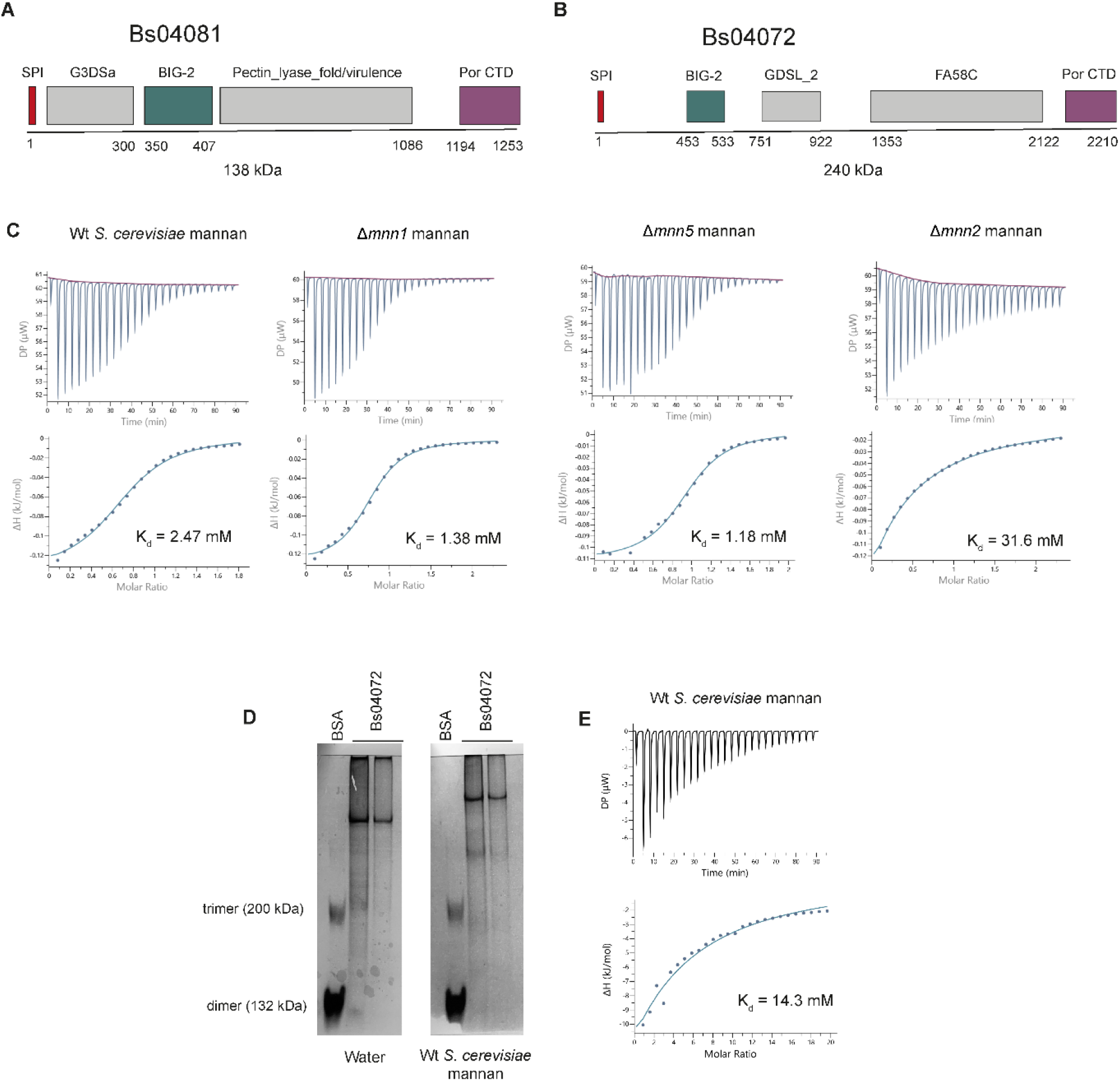
a and b) Domain organisation of Bs04081 and Bs04072. Potential binding Big-2 domain is shown in teal, T9SS C-terminal sequence in purple, SPI is red, other domains are in grey. c) Binding of Bs04081 to mannans from WT (0.16% w/v), Δmnn1 (0.5%), Δmnn5 (0.25%), Δmnn2 (0.5%) *S. cerevisiae* strains assessed by ITC. Mannans were titrated into 20-25 μM Bs04081. The stoichiometry was fitted to a one-site binding model with n of 1, to determine Kd. d) Binding of Bs04072 to 0.25% w/v mannan from WT *S. cerevisiae* analysed by NATIVE-PAGE. 5 μg of protein was loaded onto affinity gels containing 0.1% yeast mannan or water as control, BSA was used as a non-binding control. e) ITC of binding of Bs04072 to mannan from WT *S. cerevisiae*. Mannan was titrated into 20 μM Bs04081.

### *B. salyersiae* manno-oligosaccharides promote growth of other gut *Bacteroides*

We next investigated whether the oligosaccharides generated by *B. salyersiae* would enable the growth of other poor mannan degrading gut *Bacteroides*, in contrast to the ‘selfish’ mannan degradation by *B. thetaiotaomicron* (41). To address this, we cultured 19 different *Bacteroides* species on sterilised conditioned media derived from *B. salyersiae* mannan cultures. Several species that are unable to degrade polymeric mannan were able to grow on this conditioned media; *B. xylanisolvens* DSM 1836 and *NLAE-zI* p352, *B. ovatus* ATCC 8483 and CL03T12C18, as well as *B. finegoldii* DSM 17565 and CL09T03C10 (**Fig. 7a** and **Fig. S5**). In contrast, conditioned media derived from *B. thetaiotaomicron* did not allow growth of poor mannan degraders (**Fig. 7b**), consistent with previously published data (41). Analysis of the cell-free media post bacterial growth revealed that extracellular mannose was consumed by the species whose growth was not promoted by the conditioned media, whereas species that thrived in conditioned media did not deplete the mannose (**Fig S5**). This suggests that manno-oligosaccharides and not extracellular mannose serve as a nutrient source during crossfeeding.

**Figure 7.**
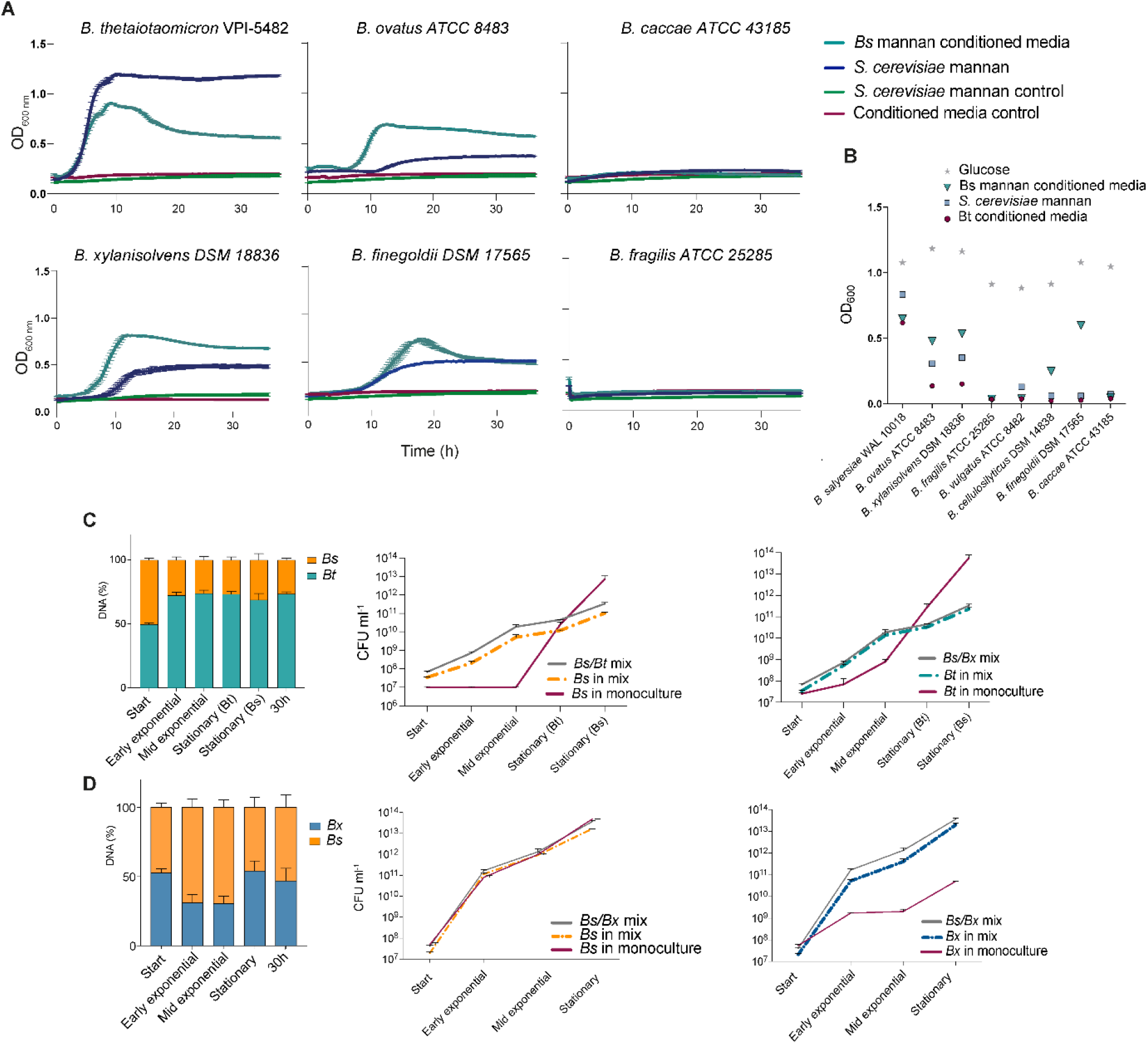
Sharing of yeast manno-oligosaccharides between gut *Bacteroides*. a) Growth of *B. thetaiotaomicron* VPI-5482; *B. xylanisolvens* DSM 18836; *B. ovatus* ATCC 8343; *B. finegoldii* DSM 17565; *B. caccae* ATCC 43185; *B. fragilis* ATCC 25285 on 10 mg ml^-1^ yeast mannan in defined medium (blue trace) or *B. salyersiae* derived conditioned media, containing digested yeast mannan (teal trace), mannan and conditioned media without bacterial growth are shown in green and maroon traces, respectively. Growth curves are averages from triplicates, and are a representative growth from at least 3 independent experiments. b) Maximum OD_600nm_ achieved by indicated strains after 28 h growth on 10 mg ml^-1^ yeast mannan, *B. salyersiae* conditioned media (green triangle) or *B. theta* conditioned media (maroon circle); growth in defined media containing 10 mg ml-1 glucose is shown with grey star. c) Competition assays between *B. salyersiae* and *B. theta* VPI-5482 d) cross-feeding assays between *B. salyersiae* and *B. xylanisolvens* DSM 18836. Bs/Bt or Bs/Bx were co-cultured in defined medium contaning yeast mannan, monocultures were set up alongside to evaluate growth. Total CFU ml^-1^ was determined from colony counts and proportions of each species was determined by qPCR from genomic DNA using marker genes for identification. Error bars are SD from 3 biological replicates, data set is representative of at least 2 independent experiments.

*B. thetaiotaomicron* outcompetes *B. xylanisolvens NLAE-zI* p352 in a co-culture on yeast mannan, but when mannose is used as a sole carbon source, *B. xylanisolvens* outcompetes *B. thetaiotaomicron* (41). To investigate whether *B. salyersiae* would share yeast mannan, we performed a set of inter-species growth assays, where *B. salyersiae* was co-cultured on either yeast mannan or mannose with a poor mannan degrader or *B. thetaiotaomicron*, a ‘selfish’ mannan degrader. This revealed that *B. salyersiae* was able to share mannose with both *B. xylanisolvens* and *B. thetaiotaomicron* (**Fig. S6**). In mannan co-cultures, *B. salyersiae* promoted growth of *B. xylanisolvens* and was able to survive competition with *B. thetaiotaomicron* (**Fig. 7b and c**). In fact, during growth with *B. thetaiotaomicron, B. salyersiae* was able to grow more rapidly in early time points, avoiding the long lag phase seen in *B. salyersiae* monoculture. Analysis of cell free supernatant taken during both co-cultures revealed that the oligosaccharides were released into the media (**Fig. S6**), a fraction of which were still present at stationary phase, suggesting that these could potentially be used by other species such *B. ovatus* and *B. finegoldii*.

## Discussion

Here we demonstrate that another member of the human gut microbiota, *Bacteroides salyersiae*, contributes to the degradation of a polysaccharide component of the cell wall of fungi. In contrast to the previously characterised ‘selfish’ mechanism of yeast α-mannan utilisation in *B. thetaiotaomicron* (41), we show that *B. salyersiae* deploys a ‘sharing’ strategy, where branched manno-oligosaccharides are released at the cell surface and then used as ‘public goods’ by other non-mannan degrading *Bacteroides* species, such as *B. xylanisolvens, B. ovatus*, and *B. finegoldii*.

Several studies have demonstrated that the members of the gut microbiota often cooperate to degrade complex carbohydrates (23, 40, 54, 55), however some display a competitive ‘selfish’ phenotype (39). Both *B. ovatus* ATCC 8483 and *B. xylanisolvens NLAE-zI* p352 encode PULs which share structural synteny with the mannan PULs in *B. thetaiotaomicron* (41). It was previously shown that the selfish mannan utilisation by *B. thetaiotaomicron* does not allow for growth of other species, despite the presence of mannan degrading enzymes in these competitors (41). Our competition and cross-feeding assays indicate that *B. salyersiae* is able to persist in co-culture with *B. thetaiotaomicron* and favours the growth of poor mannan degraders. *B. salyersiae* uses large, multimodular enzymes and binding proteins to enable extracellular degradation of mannan.

Biochemical characterisation of this alternative mechanism revealed that *B. salyersiae* uses a single PUL and an orphan CAZyme cluster to orchestrate mannan breakdown. Further analysis showed that many enzymes as well as binding proteins in the main PUL contain C-terminal signals, which direct them to the T9SS. Comparative proteomics confirmed that the proteins essential for the assembly of the T9SS were not upregulated but detected in *B. salyersiae* during growth on yeast mannan. The T9SS mediates secretion of proteases in the oral pathogen *P. gingivalis* and gliding motility proteins in the environmental Bacteroidetes *F. johnsonii* (8) It has also been shown to play a central role in chitin and cellulose metabolism in *F. johnsonii* and *C. hutchinsonii* (16, 56), respectively, but has not been described in human gut *Bacteroides* before. Our bioinformatics analysis revealed that, in addition to *B. salyersiae*, the proteins containing C-terminal T9SS signals are present in a small subset of gut *Bacteroides*, which include *B. nordii, B. intestinalis*, and *B. cellulosilyticus. B. salyersiae* possesses around 100 proteins that contain terminal T9SS signals. A small proportion of these proteins are predicted to be carbohydrate active enzymes but the function of the majority of them is unknown, indicating that, in addition to glycan degradation, T9SS contributes to multiple other processes in *B. salyersiae*.

The PUL orchestrating mannan degradation in *B. salyersiae* includes two α-mannan specific binding proteins both of which contain T9SS C-terminal motifs. Previous studies have shown that T9SS-directed proteins can be either attached to LPS at the cell surface by a sortase (13, 57) or solubly into the extracellular milieu (15). Our attempts to detect secreted enzymes or other proteins in the supernatant after growth were unsuccessful – concentrated supernatant had no detectable mannanase or mannosidase activity, and those proteins that were identified through MS of SDS-PAGE bands are likely associated with OMVs. It is likely therefore, that *B. salyersiae* follows the *P. gingivalis* paradigm of conjugating T9 secreted proteins to the surface on anionic-LPS, and T9SS cargo proteins are enriched in released OMVs (58). This is perhaps more appropriate for gut-resident Bacteroidetes, as rapid transit times and nutrient turnover means that any products of distal secreted enzymes would likely be lost to competitor species. Similarly, *Bacteroides* surface glycan binding proteins tend to be lipoproteins anchored in the outer membrane such as SusE and F from the canonical starch PUL (34, 59), and those characterised from the heparin (28) and xyloglucan PULs (60) which aid in capture of polysaccharide or oligosaccharide fragments. The T9SS secreted proteins Bs04081 and Bs04072 contain multiple predicted domains including a Big-2 domain (IPR003343, Pfam 02368), which contributes to virulence of pathogenic *E. coli*, *Citrobacte*r and *Yersinia sp*. by promoting adhesion to epithelial surfaces (61). More detailed biochemical studies will be required, however, to establish the precise binding domains and specificities of these proteins. It appears that the two central mannan-degrading enzymes, exo-acting GH38 (Bs04085) and endo-acting GH76 (Bs04077) are directed to the T9SS. Our biochemical analysis shows that contrast to the GH38 found in *B. thetaiotaomicron*, Bs04085 acts as a highly efficient broad-acting exo-α-mannosidase that matches the large amount of mannose released in the whole-cell assays. Presumably due to cellular toxicity, we were unable to clone or produce recombinant Bs04077 to test its activity. *B. salyersiae* is resistant to many of the antibiotics used in *Bacteroides* genetic manipulations, and is already able to grow on inulin, so we could not to use the elegant gain-of-function system developed by Garcia-Bayona et al (62). Unfortunately, we were therefore unable to biochemically or genetically confirm that the T9SS secreted GH76 Bs04077 is the key surface enzyme which produces the large branched oligosaccharides detected in growth and whole cell assays. However, the alternative GH76 in the *B. salyersiae* mannan PUL does not release the same pattern of oligosaccharides, though it is predicted by SignalP to be a lipoprotein and could therefore also be located on the outer surface of the cell. The majority of characterised GH76 are unable to hydrolyse highly branched mannan, and instead prefer the linear α-1,6 backbone (63). The additional ricin-lectin domain and CBM6 found in Bs04077 might enable it to target more decorated mannan structures, enabling the release of the branched oligosaccharides. One important feature of the T9SS is the ability to translocate extremely large (100-650 kDa) proteins across the outer membrane (44). The size of this multimodular enzyme (~145 kDa) is much larger than typical lipoproteins and therefore likely requires the T9SS to translocate across the outer membrane.

The branched oligos we predict are generated by Bs40477 are transported into the periplasm through the SusCD pair, where, in contrast to *B. thetaiotaomicron*, there are no other endo-mannanases, and the oligosaccharide breakdown must be achieved by exo-mannosidases from the GH92, GH38 and GH125 families.

*B. salyersiae* is not a well characterised gut *Bacteroides* and to the best of our knowledge this is the first study which describes the contribution of this bacterium to glycan metabolism. Mannans are widely recognised by a variety of receptors on immune cells and the epithelial lining (64), therefore it is possible that the enzymatic degradation of fungal cell wall would lead to altered recognition and immune response. Furthermore, fungal overgrowth is associated with Crohn’s disease as well as the abundance of Anti-Saccharomyces Cerevisiae Antibodies (ASCA) which recognise man-α-1,3-man-α-1,2 capped manno-oligosaccharides (65, 66). Our exo-mannosidase degradation assay demonstrated that at least a proportion of oligosaccharides released by *B. salyersiae* from yeast mannan retains such decoration, suggesting that this bacterium could potentially contribute to the pathogenesis of inflammatory bowel diseases. A previous multi-omics study which mapped cytokine profiles with microbial species in the human gut identified that the presence of *B. salyersiae* dampens the IFN-γ response elicited by *C. albicans* hyphae (67). *B. thetaiotaomicron* was previously shown to possess an enzyme which targets β-mannosidic linkage found in *C. albicans* mannan (68), suggesting that commensal bacteria potentially can cooperate to degrade mannan of pathogenic fungi and control their physiology. However, whether such interaction occurs *in vivo* and its role in disease states is unknown.

## Materials and Methods

### Polysaccharide sources

Wild-type mannan from *Saccharomyces cerevisiae* was purchased from Sigma. *S. cerevisiae* mannan variants were purified from *Δmnn2, Δmnn5*, and *Δmnn1* mutants (42). Cultures were grown to stationary phase, pelleted, and mannans were precipitated by Fehling reagent and washed with Methanol/Acetic acid as described by Kocourek and Ballou (69). Extracted mannans were dialysed into water and freeze-dried.

### Bacterial strains and growth conditions

The strains of *Bacteroides* spp. used in this study were as follows:

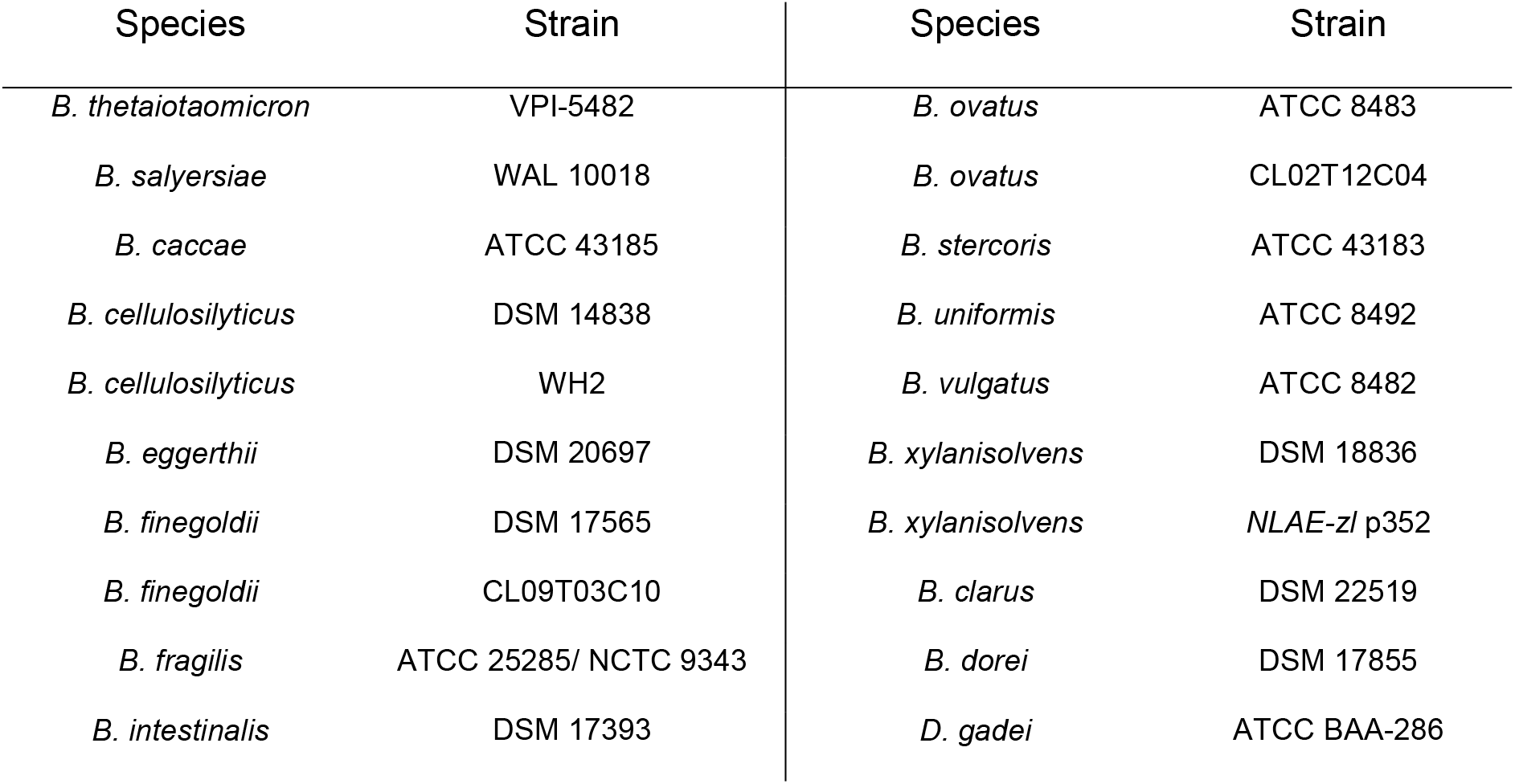

All strains were routinely grown in TYG (tryptone, yeast extract, glucose) medium supplemented with hematin for 16 h at 37 °C under anaerobic conditions, in a Whitley A35 workstation (Don Whitley). Overnight cultures were then sub-cultured into defined medium as described previously (70) and 1% w/v mannan from *S. cerevisiae* as a sole carbon source. Growth assays were performed in triplicates in a 96-well plate using a Biotek Epoch plate reader. All growth curves presented in this study are representative of at least 3 independent experiments. For conditioned media experiments, *B. salyersiae* or *B. thetaiotaomicron* were grown in defined medium with 15 mg ml^-1^ yeast mannan as a sole carbon source to mid-exponential phase (OD_600_= 0.6-0.8), supernatant was collected and autoclaved. Prior to use, conditioned media was filter sterilised and diluted with 2X defined media without a carbon source to replenish nutrients. This resulted in approximately 7.5 mg ml^-1^ of digested yeast mannan.

### *Bacteroides* co-culture experiments

Active overnight cultures of *B. thetaiotaomicron* VPI-5482, *B. salyersiae* WAL-1008, and *B. xylanisolvens* DSM 18836 were inoculated at 5 % v/v into 10 ml of defined media with 10 mg ml^-1^ glucose and grown to mid exponential phase (OD_600_ = 0.7). Cells were pelleted by centrifugation, washed three times with sterile PBS, and resuspended in 10 ml of PBS. Cells were inoculated at 5% v/v ratio into fresh defined media containing 10 mg ml^-1^ yeast mannan. Respective monocultures on 10 mg ml^-1^ yeast mannan were also included. All cultures were set up in triplicates. Co-culture samples (1 ml) were collected at the point of inoculation, early exponential (OD_600_ = 0.4), mid exponential (OD_600_ = 0.7-0.8), early stationary (OD_600_ = 2.0), and late stationary growth phases. For the *B. salyersiae/B. thetaiotaomicron* co-culture stationary phases were collected for each species. Genomic DNA was extracted and proportion of each species in a co-culture was determined by absolute quantification qPCR using primers for *Bt3780, Bs04085, Bxy_2930* to identify *B. thetaiotaomicron, B. salyersiae*, and *B.xylanisolvens*, respectively. Total gDNA concentration was measured using Qubit Fluorometer (Thermo Fisher) and diluted to 10 ng μl^-1^, concentration of species-specific DNA was extrapolated from the standard curve. qPCR was performed using Luna Universal Mastermix (NEB), denaturation/annealing/elongation was performed for 50 cycles, Cq was read by exciting samples at 497 nm and measuring light emission at 520 nm. Samples were also serially diluted, plated on TYG plates, and incubated anaerobically for 2 days to determine CFU ml^-1^ counts, CFU ml^-1^ of each species in the co-culture was extrapolated from total counts using qPCR data.

### Enzymatic assays with whole cells

*B. salyersiae* was grown in 5 ml of defined media containing 10 mg ml^-1^ yeast mannan to OD_600_=0.8. The cells were then collected by centrifugation at 5000 rpm for 10 min and washed with sterile PBS (pH 7.0) three times and resuspended in 1.75 ml of PBS. Cells (250 μl) were then incubated with 10 mg ml^-1^ yeast mannan to investigate enzymatic activity at the cell surface. As a control, surface enzymes were removed with Proteinase K (2 mg ml^-1^) treatment for 2 hours at 37 °C, cells (500 μl) were washed 3 times with sterile PBS to remove residual protease and resuspended in 500 μl of PBS. A proportion of these were incubated with 10 mg ml^-1^ yeast mannan. To demonstrate intracellular activity, both intact and Proteinase K treated cells were lysed with BugBuster (Merk Millipore) and assayed against 10 mg ml^-1^ yeast mannan. The supernatant without cells was also tested against yeast mannan to demonstrate that the enzymes were not present in the extracellular milieu. Cells resuspended in PBS without a carbon source were also included as control. All assays were performed in parallel at 37 °C and 50 μl aliquots were taken at t= 0; 0.2; 0.5; 1, 2, 3, 4, and 16 h.

### Thin layer chromatography

Aliquots of 3 to 9 μl were spotted onto silica coated plates and resolved in butanol/acetic acid/water (2:1:1), sugars were visualised with orcinol/sulphuric acid stain.

### Cloning, expression, and purification of recombinant proteins

DNA encoding the genes of interest, excluding the signal peptide, were amplified by PCR with appropriate primers from *B. salyersiae* genomic DNA and cloned into pET28-a vector using NcoI/XhoI or NheI/XhoI restriction sites. Recombinant proteins contained His6-tags at the C-terminus, except for BS_04078^GH92^, which had an N-terminal His-tag. Genes from *B. thetaiotaomicron* were cloned in a previous study (41, 43). To produce recombinant proteins, plasmids were transformed into TUNER *E. coli* strains and grown in of LB broth with 10 μg ml^-1^ kanamycin to mid-exponential phase at 37 °C. The cultures were then cooled and protein expression was induced with 0.2 mM isopropyl-β-D-galactopyranoside (IPTG) at 16 °C with shaking at 180 rpm for 16 h. Recombinant proteins were then purified from cell-free extracts by immobilised metal affinity chromatography, using a cobalt-based matrix (Talon, Clontech). The size and purity were assessed with SDS-PAGE, concentrations were calculated using absorbance at 280 nm (NanoDrop, Thermo Scientific) and extinction coefficients.

### Assays with recombinant enzymes

Enzymatic activity was tested at 1 μM enzyme against 4 mg ml^-1^ yeast mannan or 1 mM disaccharide, where appropriate. All reactions were performed in 50 mM MOPS containing 2 mM CaCl2 buffer at 37C overnight, unless stated otherwise. Continuous mannose release was determined using a linked enzyme assay system (D-glucose/D-fructose/D-mannose detection kit, Megazyme), as described in (41) to assess kinetic parameters and perform end point assays with α-mannosidases. Mannose concentration was calculated using extinction coefficient of NADPH (6230 M^-1^ cm^-1^) at 340 nm.

### Binding assays

Binding affinity was assessed with polyacrylamide gel electrophoresis. Gels, containing 0.1% (w/v) yeast mannan or water as control, were loaded with 5 μg of protein and run at 10 mA per gel. Gels with and without ligand were run in the same tank. BSA (15 μg) was used as a non-interacting control.

The binding was further quantified by Isothermal Titration Calorimetry. In brief, titrations were performed at 25 °C and 307 rpm with 27 injections at regular intervals of 10 μl of ligand into the cell containing 25 μM of protein. The concentration of ligand was adjusted depending on the conditions to obtain a titration curve. The curve was analysed with non-linear regression and was fitted to a model where number of binding sites was equal to 1, using Microcal PEAQ-ITC software v1.30 (Malvern Panalytical).

### OMV purification and Transmission Electron Microscopy

*B. salyersiae* was grown in 50 ml of defined medium containing either 10 mg ml^-1^ mannan from *S. cerevisiae* or glucose anaerobically at 37 °C for 16 hours. Cultures were centrifuged at 10,000 rpm for 20 minutes, supernatant was collected, and filter sterilised. The filtrate was then ultracentrifuged 100,000 g for 2 hours (Optima L-90K, Beckman Coulter). The vesicle pellet was resuspended in 100 μl of sterile PBS.

Samples were then soaked onto carbon-coated copper grids, negatively stained with 2 % aqueous uranyl acetate for 15 mins and analysed with Transmission Electron Microscope (Hitachi HT7800 120 kV). Pictures were taken at 100 kV accelerating voltage with 20,000 x magnification. The diameters of OMVs were measured in Fiji.

### Analysis of in-gel proteins by Mass Spectrometry

Mass Spectrometry analysis of in-gel proteins was performed by the Metabolomics and Proteomics facility, Department of Biology (University of York, UK), as a paid service. In brief, *B. salyersiae* was cultured in defined medium with 10 mg ml^-1^ yeast mannan, aliquots were collected hourly. Cell free supernatant was concentrated 5 times with Vivaspin® 20 concentrator tubes with 30 kDa cut off, 20 μl was analysed with SDS-PAGE gel. Desired protein bands were excised from the gel and sent off for analysis. Proteins were trypsin digested, the resulting peptides were analysed with Matrix Assisted Laser Desorption Ionization Tandem Time-of-Flight (MALDI-TOF) mass spectrometry. Mass spectral data was searched against *B. salyersiae* CL02T12C01 genome available in the NCBI database using Mascot search program. Threshold for the associated expect value was set to 0.05, fragment mass tolerance was set at 0.5 Da, a minimum of 2 matched peptides was required for protein identification.

### Comparative proteomics

#### Sample preparation

*B. salyersiae* cells were suspended in 5% sodium dodecyl sulfate (SDS) in 50 mM triethylammonium bicarbonate (TEAB) pH 7.5. The samples were subsequently sonicated using an ultrasonic homogenizer (Hielscher) for 1 minute. The whole-cell lysate was centrifuged at 10,000 x g for 5 minutes to remove cellular debris. Protein concentration was determined using a bicinchoninic acid (BCA) protein assay (Thermo Scientific). Proteins (30 μg) were reduced by incubation with 20 mM tris(2-carboxyethyl)phosphine for 15 minutes at 47 °C, and subsequently alkylated with 20 mM iodoacetamide for 30 minutes at room temperature in the dark. Proteomic sample preparation was performed using the suspension trapping (S-Trap) sample preparation method (71, 72), as recommended by the supplier (ProtiFi™, Huntington NY). Briefly, 5 μl of 12% phosphoric acid was added to each sample, followed by the addition of 330 μl S-Trap binding buffer (90% methanol in 100 mM TEAB pH 7.1). The acidified samples were added, separately, to S-Trap micro-spin columns and centrifuged at 4,000 xg for 1 minute until all the solution has passed through the filter. Each S-Trap micro-spin column was washed with 150 μl S-trap binding buffer by centrifugation at 4,000 xg for 1 minute. This process was repeated for a total of five washes. Twenty-five μl of 50 mM TEAB containing trypsin (1:10 ratio of trypsin:protein) was added to each sample, followed by proteolytic digestion for 2 hours at 47 °C using a thermomixer (Eppendorf). Peptides were eluted with 50 mM TEAB pH 8.0 and centrifugation at 1,000 xg for 1 minute. Elution steps were repeated using 0.2% formic acid and 0.2% formic acid in 50% acetonitrile, respectively. The three eluates from each sample were combined and dried using a speed-vac before storage at −80°C.

#### Mass spectrometry

Peptides were dissolved in 2% acetonitrile containing 0.1% trifluoroacetic acid, and each sample was independently analysed on an Orbitrap Fusion Lumos Tribrid mass spectrometer (Thermo Fisher Scientific), connected to an UltiMate 3000 RSLCnano System (Thermo Fisher Scientific). Peptides (1 μg) were injected on a PepMap 100 C18 LC trap column (300 μm ID × 5 mm, 5 μm, 100 Å) followed by separation on an EASY-Spray nanoLC C18 column (75 μm ID × 50 cm, 2 μm, 100 Å) at a flow rate of 250 nl min^-1^. Solvent A was water containing 0.1% formic acid, and solvent B was 80% acetonitrile containing 0.1% formic acid. The gradient used for analysis of proteome samples was as follows: solvent B was maintained at 2% for 5 min, followed by an increase from 2 to 35% B in 120 min, 35-90% B in 0.5 min, maintained at 90% B for 4 min, followed by a decrease to 3% in 0.5 min and equilibration at 2% for 10 min. The Orbitrap Fusion Lumos was operated in positive-ion data-dependent mode. The precursor ion scan (full scan) was performed in the Orbitrap in the range of 400-1,600 m/z with a resolution of 120,000 at 200 m/z, an automatic gain control (AGC) target of 4 × 10^5^ and an ion injection time of 50 ms. MS/MS spectra were acquired in the linear ion trap (IT) using Rapid scan mode after high-energy collisional dissociation (HCD) fragmentation. An HCD collision energy of 30% was used, the AGC target was set to 1 × 10^4^ and dynamic injection time mode was allowed. The number of MS/MS events between full scans was determined on-the-fly to maintain a 3 s fixed duty cycle. Dynamic exclusion of ions within a ± 10 ppm m/z window was implemented using a 35 s exclusion duration. An electrospray voltage of 2.0 kV and capillary temperature of 275°C, with no sheath and auxiliary gas flow, was used.

#### Mass spectrometry data analysis

All spectra were analysed using MaxQuant 1.6.14.0 (73), and searched against the Uniprot *Bacteroides salyersiae* CL02T12C01 proteome database (UP000005150) downloaded on 01 October 2020. Peak list generation was performed within MaxQuant and searches were performed using default parameters and the built-in Andromeda search engine (74). The enzyme specificity was set to consider fully tryptic peptides, and two missed cleavages were allowed. Oxidation of methionine, N-terminal acetylation and deamidation of asparagine and glutamine were allowed as variable modifications. Carbamidomethylation of cysteine was allowed as a fixed modification. A protein and peptide false discovery rate (FDR) of less than 1% was employed in MaxQuant. Proteins that contained similar peptides and that could not be differentiated on the basis of MS/MS analysis alone were grouped to satisfy the principles of parsimony. Statistical analysis was performed in the R statistical programming language using the MSstats package (75).). Since the *B. salyersiae* DSM 18765/WAL10018 strain is not in the UniProt data base, identified proteins were matched from *B. salyersiae* CL02T12C01 to *B. salyersiae* DSM18765/WAL10018 using the ‘Genome gene best homologs’ tool at https://img.jgi.doe.gov. A full list of equivalent locus tags can be found in **Table S4.** Percentage identity across the mannan PUL and cazyme cluster genes ranged from 99.1-100%.

#### Bioinformatics

Protein sequences were retrieved from the Integrated Microbial Genomes (IMG) database (76). Sequence alignments and matrix identities were perfomed in Clustal Omega (77). The presence and nature of signal peptides was determined in SignalP 5.0 (78). Modular organisation of proteins was assessed using InterPro (79). Glycoside hydrolase families and PUL boundaries were identified in the Cazy database (http://www.cazy.org) and PULDB (45).

## Acknowledgements

We thank Dr David Bolam, Dr Fiona Cuskin and Professor Janet Quinn (Newcastle University) for advice and discussions. We would like to thank Tracey Davey, EM Unit, Newcastle University, for expert technical assistance. The TEM equipment in the EM facility was funded by BBSRC grant BB/R013942/1. This work was funded by a Newcastle University Faculty PhD studentship awarded to ECL.

## Author contributions

EB and ECL designed the research, analysed data and wrote the manuscript. EB carried out most of the experimental work. CM cloned enzymes and purified proteins, CC cloned and characterised BS04078 and performed assays with BS04085. TH and MT collected and analysed Mass Spectrometry data.

## Notes

### Competing Interest Statement

The authors have declared no competing interest.

