## Supplemental Figs 1-6 for "The Type 9 Secretion System enables sharing of fungal mannan by human gut *Bacteroides*"

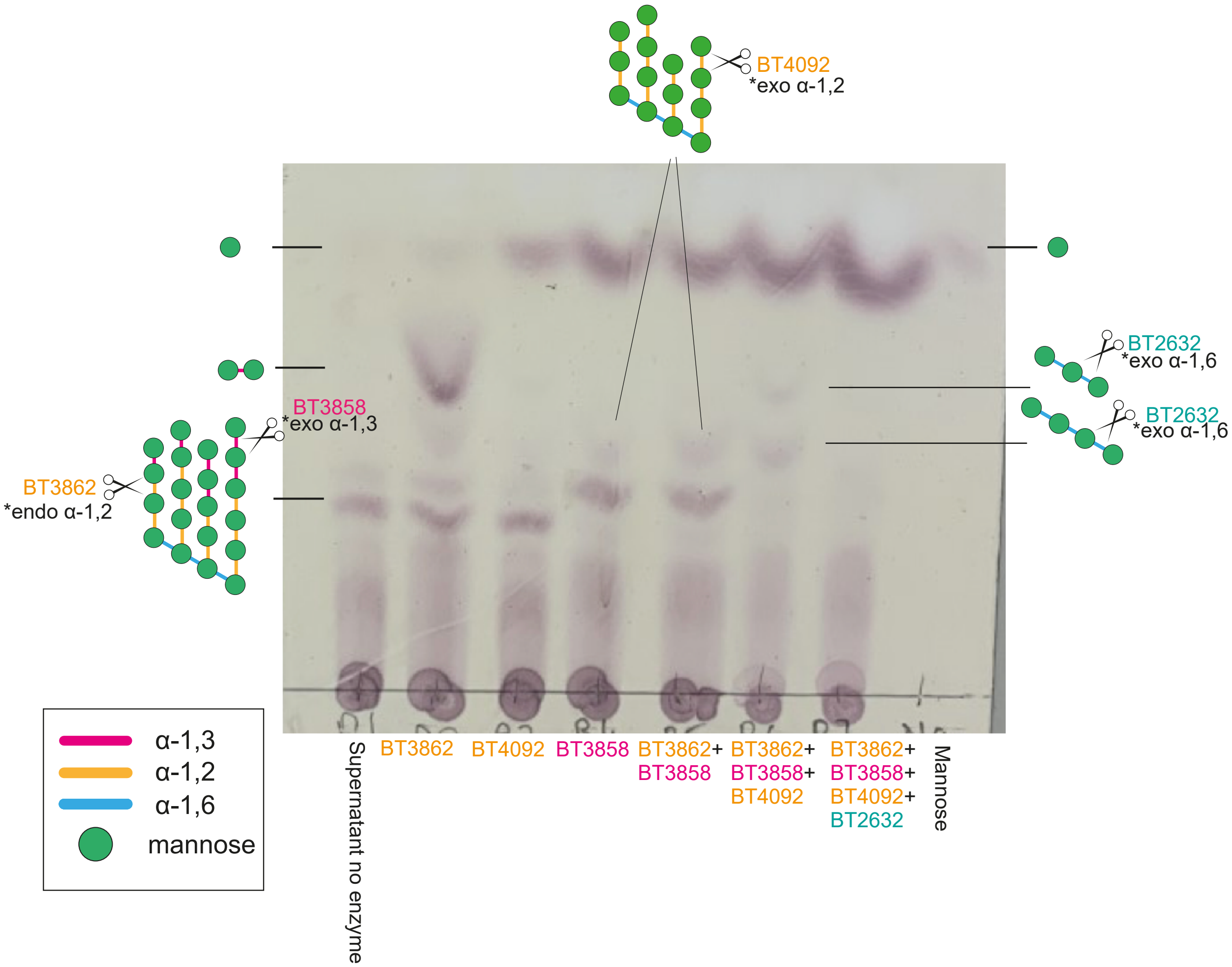


Supplementary Figure 1. *B. salyersiae* releases branched manno-oligosaccharides from yeast α-mannan. *B. salyersiae* supernatant collected from the stationary phase was sequentially degraded by previously characterised α-mannosidases from *B. thetaiotaomicron*: BT3862, endo α-1,2-mannosidase, releases α-1,3 mannobiose; BT3858 and BT4092 - exo-α-1,3 and exo-1,2 mannosidases, respectively, act on branched mannan; BT2632 - exo-α-1,6 mannosidase breaks down linear α-1,6 mannooligosaccharides.

**A**

```
Bs04091 GH97      -----------------------------DFLDPGKK-YMATV-YADAADTDYENNQQAY   29
Bs04089 GH125     -------------------------------------------------------DEEIK   5
Bs04085 GH38      ---------------------------------------------------------NVY   3
PG0026            ---------------------------------------------------------EIY   3
Bs04078 GH92      ---------------------------SAPENSDKSR-FVQEVKYNGKN------YSKNF   26
Bs04076 GH2       ---------------------------------NGEI-AV---CTPGKA------VIKVM   17
Bs04071 unk       ----------------------------TIYSDNGGL-LI---KSHA-DVT---GYYELF   24
Bs04075 unk       -----------------------TLENIAVYSISGGI-EM---IA---SQA---VQINIY   27
Bs04081 unk       -------------------------------TSDNAI-F-----VSGMKNT---AKIALY   20
Bs04073 GH78      ----------------------------------GRI-RV---QLKNDYRA---DIVTLF   19
Bs04077 GH76      -------------------------------IINNRL-YI---DVPETA------QVQVC   19
Bs04072 unk       -----------------------------PVSRGTKV-H-----LNGNG------HVSVC   19
Bs04074 GH3       -----------------------------VVEAGESV-NI---SFGIVA------KADVY   21
Bs01537 GH38      --------------CFLQGKNHPDEEWKTLDRIDGNSRNKVERQLPQPAQA---------   37
Bs04086 GH2       VNDLYLSPNPTKGYCSIEGTNSELINEVRIFSLSGKT-----------------------   37
Bs04088 GH3       ---------------------KELKQFKKVFLKAGES-----------------------   16
Bs04090 GH76      ----YKQD----------NNKKYLDAFNRSLT----------------------------   18

                    Motif B                                       Motif D
Bs04091 GH97      TIRKMVVTNKS--------------------------KFKQSVSIGSGFAISLFEITDNK   63
Bs04089 GH125     DCVKMLIDTDAGTGFIHESFHKDDPANFTRAWFAWQNT-------LFGE--LILKLVNEG   56
Bs04085 GH38      DCIGKVIDNR---VY---PVSKSGLQ-----EFIWDSH-----DINEGI--YLYTISAMK   45
PG0026            DFTGRLVNSLPVKTY---SSSYGEPI-----EIKWDLTSKYGVKIGNGF--YLYRCVVNS   53
Bs04078 GH92      LVHSSLLNGADIQFR---MGD----------------KPNKKRGINKAD--YPYSLSNEK   65
Bs04076 GH2       NVGGYILA--DYQS----EG-----------------KLTIGMNYTDGV--YLVGVETEN   52
Bs04071 unk       SIAGQCLS--KGKI-----GP----------------GTTQREYLLSGT--YLVSLEIRG   59
Bs04075 unk       AIDGRLVR--SVMLN---E------------------GRNSITGIAPGL--YLVNRT---   59
Bs04081 unk       NINGQLVS--NVIT----SGE-----------------NIELPIMGKGF--YIVRIVEDE   55
Bs04073 GH78      DMSGKSIV--VKKKQ---SGN-----------------FHIGSNLHKGI--YTVSLECNK   55
Bs04077 GH76      DLTGMLIH--DEKIQ---SGV----------------SYIPMANLNGGM--YIVRIQLSN   56
Bs04072 unk       SLQGILIA--EYY-F---SAD----------------CVIPTDGLKAGI--YIVRLDNGE   55
Bs04074 GH3       NTSGVLKA--NLY------NT----------------DRIPTTGLTQGM--YVVHMCVND   55
Bs01537 GH38      RYIRLLVTN---------------------PVQDA-------EGKDARI--YEFEVYK--   65
Bs04086 GH2       E---KTFSN---------------------KVFNIDD-------MDKGI--YLIQ-AYTK   63
Bs04088 GH3       ITIKMLLSK---------------------DAFTYYNIAAKSFLKDKGD--YHIMLGFSS   53
Bs04090 GH76      --------------------------------YAWDN-----ARDDNGL--FNVDLSGVD   39

                          Motif E
Bs04091 GH97      KK------------------------   65
Bs04089 GH125     KTDLLNSIQ-----------------   65
Bs04085 GH38      DGKMIFTDTKKMIISKFVAE------   65
PG0026            PGGQTASMAKKMIVVGQ---------   70
Bs04078 GH92      --------------------------   65
Bs04076 GH2       IPRM-----ATYKVMLCR--------   65
Bs04071 unk       QKEM-----RK---------------   65
Bs04075 unk       ----------KVIVTD----------   65
Bs04081 unk       MTTT-----VKVVVK-----------   65
Bs04073 GH78      EVYS-----QKVIIP-----------   65
Bs04077 GH76      RTIS-----RKFIK------------   65
Bs04072 unk       EIKT-----GKLIVK-----------   65
Bs04074 GH3       KKIN-----GKFIVK-----------   65
Bs01537 GH38      --------------------------   65
Bs04086 GH2       -AG-----------------------   65
Bs04088 GH3       -HDI--------CVQKNVRIL-----   65
Bs04090 GH76      KDGKKWLLTQAAIVEMYARLAQIKDL   65
```

Supplementary Figure 2. Proteins involved in mannan degradation possess T9SS targeting CTDs. Sequence alignment of the C-terminal sections of proteins upregulated during growth on mannan. T9SS motifs are highlighted, with strictly conserved residues in green, PG0026, the experimentally verified T9SS secreted sortase from *P. gingivalis* is included for comparison.

**A**

| | | | | |
|---|---|---|---|---|
| 1: BT4072GH38 | 100.00 | 21.48 | 21.09 | 20.56 |
| 2: Bs04085GH38 | 21.48 | 100.00 | 40.34 | 41.70 |
| 3: Bs01537GH38 | 21.09 | 40.34 | 100.00 | 69.42 |
| 4: BT3774GH38 | 20.56 | 41.70 | 69.42 | 100.00 |

**B**

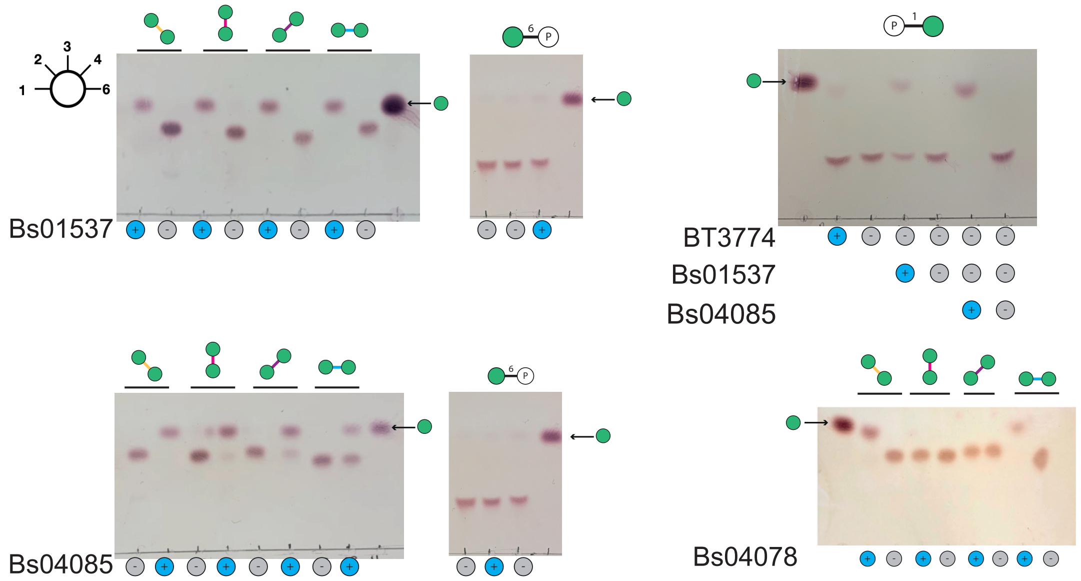


**C**

Bs01537

| Substrate | $k_{cat}/K_m$ (μM$^{-1}$min$^{-1}$) |
|---|---|
| α-1,2man2 | 2.23 |
| α-1,3man2 | 0.77 |
| α-1,4man2 | 0.35 |
| α-1,6man2 | 0.29 |

Bs04085

| Substrate | $k_{cat}/K_m$ (μM$^{-1}$min$^{-1}$) |
|---|---|
| α-1,2man2 | 7.8 |
| α-1,3man2 | 0.35 |
| α-1,4man2 | n.d |
| α-1,6man2 | n.d |

| Enzyme | Substrate | $K_m$ (mg ml$^{-1}$) | $k_{cat}$ (min$^{-1}$) | $k_{cat}/K_m$ (min$^{-1}$ mg$^{-1}$ ml) |
|---|---|---|---|---|
| Bs01537 | WT mannan | 12.04±3.2 | 15.8±3.2 | 1.31 |
| Bs01537 | Δ*mnn1* mannan | 3.2±0.4 | 86.158±3.7 | 26.92 |
| Bs01537 | Δ*mnn5* mannan | 0.777±0.2 | 71.56±4.9 | 92.69 |
| Bs04085 | WT mannan | 8.006±2.8 | 115.8±15.8 | 14.46 |
| Bs04085 | Δ*mnn1* mannan | 5.4±0.7 | 86.33±5.1 | 15.98 |
| Bs04085 | Δ*mnn5* mannan | 0.98±0.1 | 139.4±4.8 | 142.25 |

Supplementary Figure 3. a) Identity matrix of GH38s from *B. salyersiae* and *B. thetaiotaomicron* generated in Clustal omega (https://www.ebi.ac.uk/Tools/msa/clustalo/) using amino acids sequences of proteins shown. b) TLC of activity of GH38s: Bs01537 and Bs04085 and GH92: Bs04078 against α-linked mannobioses, mannose-1-phosphate, and mannose-6-phosphate. Enzymes at 1μM were incubated with 1 mM of indicated substrates in 50 mM MOPS 2 mM $CaCl_2$ overnight at 37 ºC. c) Kinetic analysis of Bs01537 and Bs04085 against α-mannobioses (left) and mannans from the wild-type and mannosyltransferase deficient *S. cerevisiae* strains (right). Reactions were performed in 40 mM MOPS 2 mM $CaCl_2$, continuous mannose release was monitored at 340 nm using Mannose detection kit. For complex polysaccharides $k_{cat}/K_m$ was determined using linear regression , for mannobiose substrates kinetics was measured below $K_m$. Data is generated from at least 3 biological replicates from 2 technical repeats, error represents SEM

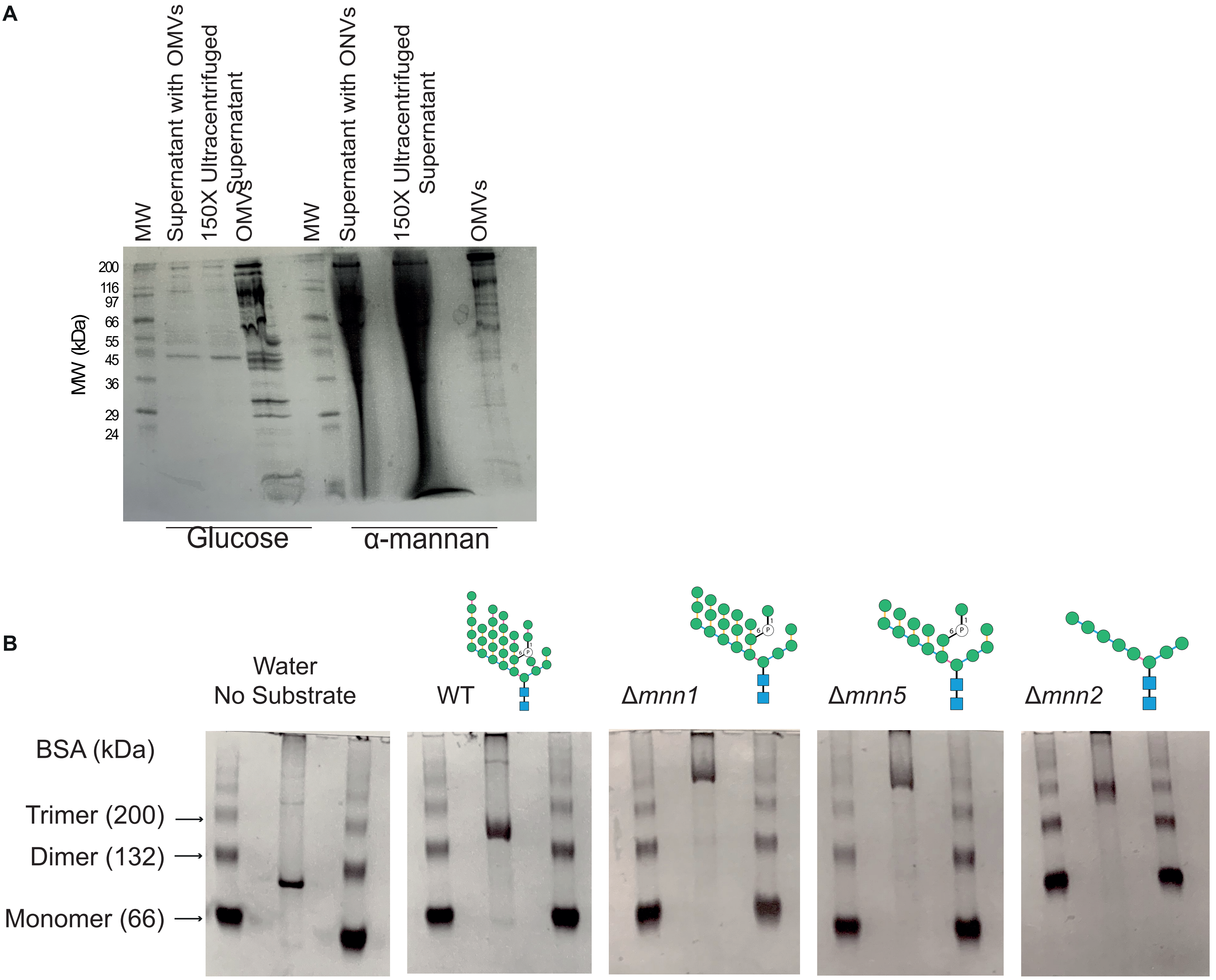


Supplementary Figure 4. a) SDS-PAGE of the supernatant and OMVs produced by *B.salyersiae* during growth on glucose or yeast mannan. OMVs were purified by ultracentifugation and resuspended in PBS, supernatant post ultracentrifugation was concentrated 150 times using a 10 kDa cut-off spin concentrator. Samples of 20 µl were loaded on the gel.b) Binding of Bs04081 to α-mannan from wild type and Δ*mnn1*, Δ*mnn5*, Δ*mnn2 S. cerevisiae*. 5 µg of protein was resolved on native PAGE affinity gel containing 0.1% (w/v) mannans or water, Bovine Serum Albumin was used as a ladder.

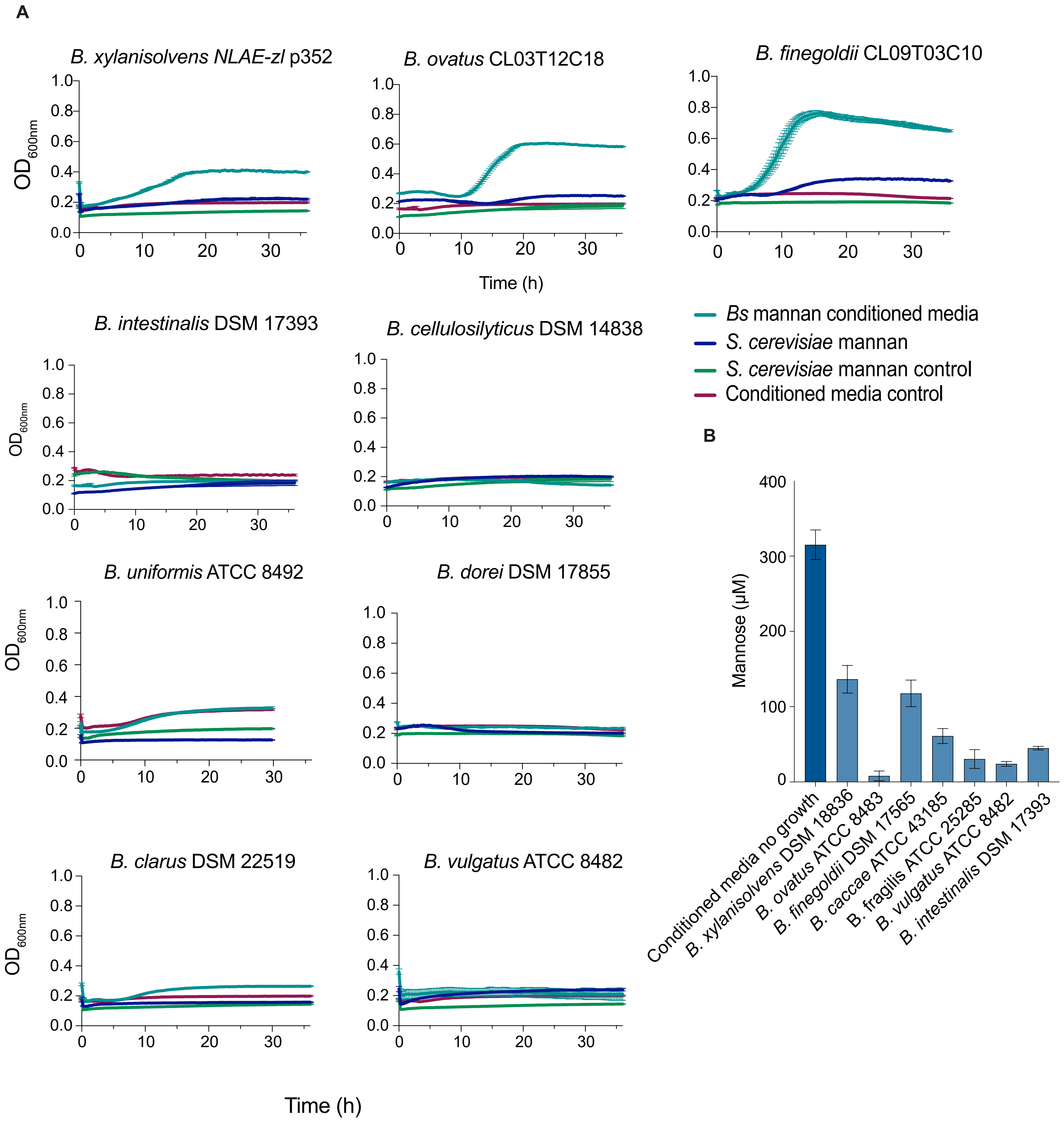


Supplementary Figure 5. a) Growth of *B. xylanisolvens* NLAE-zl p352 ; *B. ovatus* CL03T12C18; *B.finegoldii* CL09T03C10; *B. intestinalis* DSM 17393; *B. cellulosilyticus* DSM 14838; *B. uniformis* ATCC 8492; *B. dorei* DSM 17855; *B. clarus* DSM 22519; *B. vulgatus* ATCC 8482 on 10 mg $ml^{-1}$ yeast mannan in defined medium (blue trace) or *B. salyersiae* derived conditioned media, containing digested yeast mannan (teal trace), mannan and conditioned media without bacterial growth are shown in green and maroon traces, respectively. b) Mannose concentration in *B. salyersiae* derived conditioned media prior and post 38 h bacterial growth, error bars show SEM from 3 biological replicates. Dataset is representative of at least 3 independent experiments.

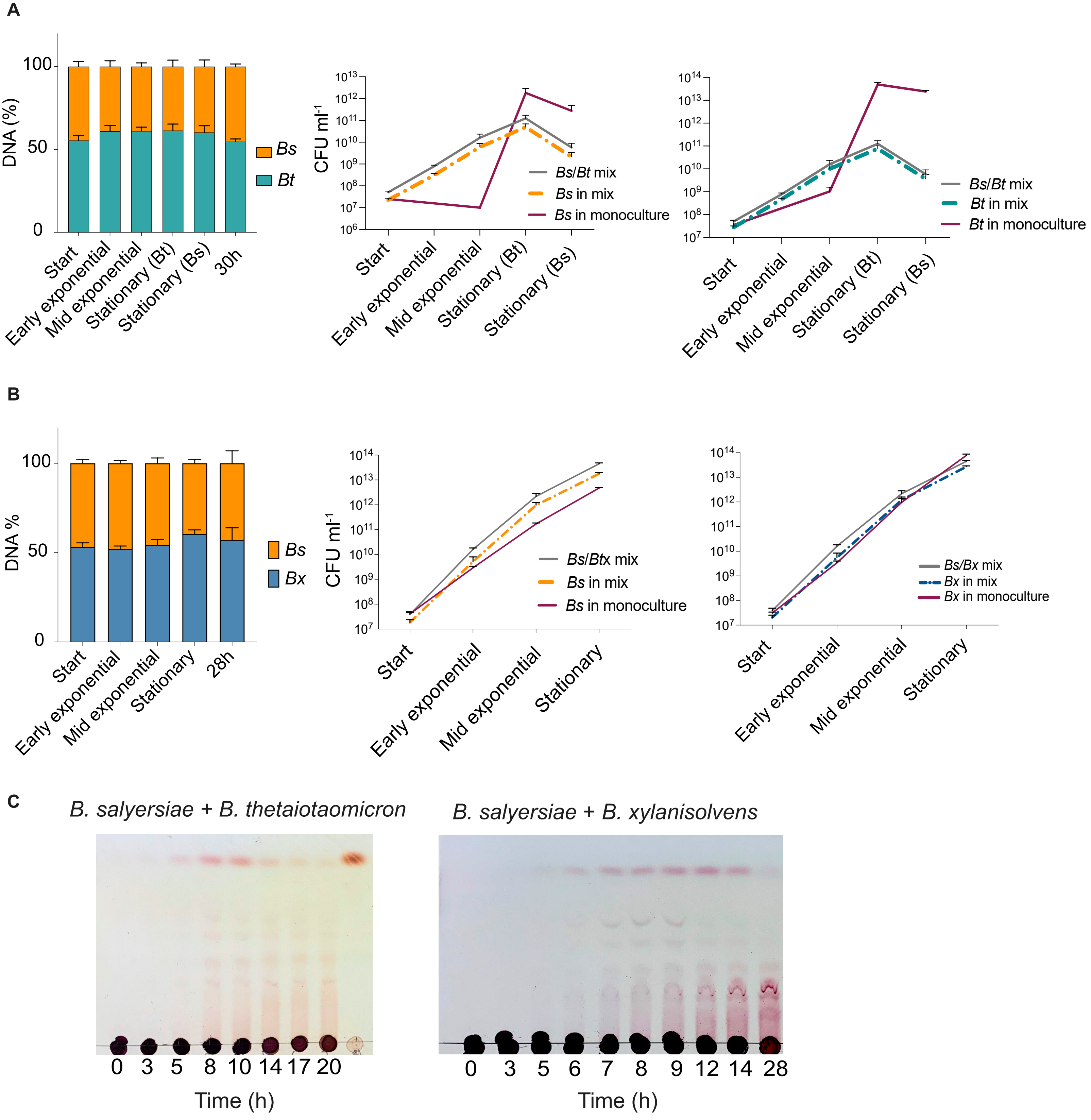


Supplementary Figure 6. Competition assays between *B. salyersiae* and *B. thetaiotaomicron* VPI-5482 (a) and *B. salyersiae* and *B. xylanisolvens* DSM 18836 (b) on mannose. Bs/Bt or Bs/Bx were co-cultured in defined medium contaning 10 mg $ml^{-1}$ mannose, monocultures were set up alongside. Total CFU $ml^{-1}$ was determined from colony counts and proportions of each species was determined by qPCR from genomic DNA using marker genes for identification. Error bars are SD from 3 biological replicates, data set is representative of at least 2 independent experiments; c) TLC of cell free supernatant from *B. thetaiotaomicron* - *B. salyersiae* or *B. salyersiae* - *B. xylanisolvens* co-cultures on yeast mannan.
